# P4HA1 and P4HA2 accept both shared and isoenzyme-specific substrate residues in type I collagen to directly and indirectly orchestrate its post-translational modification landscape

**DOI:** 10.64898/2026.10.09.757881

**Authors:** Déborah Helle, Anne Robert, Nick Ramalanjaona, Catherine Bui, Sandrine Gulberti, Jean-Baptiste Vincourt

**Author notes:** **Corresponding author:****; address:** UMR 7365, Université de Lorraine, CNRS, IMoPA, Biopole du Campus Brabois-Santé, 9, Avenue Forêt de Haye, BP20199, 54505 Vandoeuvre les Nancy Cedex France.

## Abstract

Type I collagen, the most abundant protein of vertebrate organisms, is polymerized into a macroscopic-scale fibrillar network, the detailed arrangement of which is controlled by numerous post-translational modifications (PTMs) to determine the biomechanics of connective tissues. Its most frequent post-translational modification, prolyl-4-hydroxylation (P4H), catalyzed by prolyl-4-hydroxylases (P4HAs), occupies about 100 residue positions per procollagen alpha chain and is crucial to stabilize its triple helical building block thermodynamically. Recently, the two main P4HAs, P4HA1 and P4HA2, were found to exhibit mutual substrate specificity based on the nature of the amino acid residue standing N-terminally to the considered substrate residue. Independently, the activities of both enzymes were suggested to influence lysyl residue hydroxylation and glycosylation. To clarify these findings, we performed here a proteome-, tripeptide- and amino acid-scale analysis of prolyl and lysyl modifications of collagens, both in the established ATDC5 cell line and in primary mouse skin fibroblasts after adenovirus-mediated Crispr/Cas9 invalidation of either or both *P4ha1* and/or *P4ha2* genes. We conclude that both enzymes have no more than 18 and 14 respective specific substrate positions among more than 200 potential P4H sites available on type I collagen alpha chains. Combining procollagen tryptic peptide and tripeptide analysis, we confirm that glutamate localized N-terminally to potential P4H sites determines the substrate specificity of P4HA2. P4HA1-specific sites are instead preceded by phenylalanine, lysine, arginine, asparagine or histidine. Furthermore, we evidence that P4H is not required for prolyl-3-hydroxylation at the so-called A1 site and demonstrate that it reduces lysyl glycosylation at multiple sites in a cell-type-dependent manner. These findings highlight the strategic molecular function of P4HA1 and P4HA2 within the collagen maturation machinery, for, together, steering subtle collagen post-translational modifications.

## INTRODUCTION

Type I collagen, the most abundant organic compound of the human body, is a protein assembling into a fibrillar network to determine the biomechanical properties of most connective tissues [1]. Its fibrillar assembly is finely tuned, in each tissue specifically, for the extracellular matrix to achieve functionally adapted mechanical properties. This is largely controlled by Post-Translational Modifications (PTMs) targeting prolyl and lysyl residues, the defects of which cause multiple developmental diseases [2–7] and the dysregulation of which contributes to cancer [8–10]. More specifically, oxidative deamination, hydroxylation and glycosylation of key lysyl residues are critical in establishing the valency, structure, and residue position of collagen intermolecular cross-links [11–15], which differ between tissues [12, 15, 16]. Prolyl-3-hydroxylation (P3H), another PTM targeting prolyls standing on the X position of the collagen-typical XYG amino acid tripeptide, is thought to regulate particular interactions between collagen triple helices [17–19]. It is tissue-as well as developmentally regulated [20–22], except at the so-called, C-proximal, A1 site, which in type I collagen exists only on the α1 chain [17, 19]. However, the mechanisms controlling the levels of each of these PTMs in a tissue-specific manner remain far from fully understood.

By far, the most abundant PTM of collagen is prolyl-4-hydroxylation (P4H), targeting prolyl residues occupying the Y position of the collagen-typical XYG tripeptides. It occupies about 100 residues per procollagen polypeptide [23] and is crucial to thermodynamically stabilize the procollagen triple helix assembly [24, 25], partly by promoting inter-helix hydrogen bounding [26]. Being required for collagen biosynthesis, it was long overlooked as a contributor to tissue-specific collagen PTMs. Recently, the residue positions at which type I collagen is 4-hydroxylated were found to exhibit tissue-specific profiles, tendon being particularly poorly hydroxylated at EPG tripeptides, compared to bone and skin [27]. In addition, P4HA1 and P4HA2, the two main Prolyl-4-Hydroxylases (P4HAs) in vertebrates, exhibit substrate specificity, not towards particular collagen α chains, but within the chains, towards particular residue positions [27, 28]. More specifically, based on collagen tryptic peptide analysis in mutant cells and animals, P4HA2 better targets XPG collagen tripeptides where the X residue is acidic [27, 28]. The substrate specificity of P4HA1 is more complex as it appears to target XPGs where X is either a large hydrophobic or a basic residue [28]. The substrate specificity of P4HA2 correlates to the tendon-specific P4H profile in both mouse and chicken, suggesting specialized functional consequences of P4HA2 activity [27]. Finally, several findings suggest interplays between the distinct chemical modifications of collagen [6, 19, 29, 30].

In order to better understand the detailed consequences of P4HA1 and P4HA2 activity towards chemical modifications of collagen, here, we invalidated either or the two corresponding genes, first in ATDC5 cells [27] and characterized the resulting proline and lysine modifications of collagen, at a tryptic peptide, collagen tripeptide and amino acid level. Our findings on the substrate specificity of P4HA1 differ in details from those of [28]. Also, we find that P4H does not favor P3H at the A1 site, while it limits lysine glycosylation. To avoid model-specific findings, we verified our conclusions in primary mouse skin fibroblasts.

## RESULTS

### Functional invalidation of *P4ha1, P4ha2*, or both of them in ATDC5 cells

To investigate the detailed functions of P4HA1 and P4HA2 (P4HA1&2), we took advantage of a cellular model of collagen assembly [31], already used in our previous study [27](Fig 1). It is interesting for such studies in the sense that these cells assemble collagen into a stable crosslinked network, unlike most other established cell lines [27], but their viability is not affected by the invalidation of *P4ha1* and/or *P4ha2* (data not shown). This is in contrast with the *in vivo* situation, the mouse *P4ha1* knock-out being lethal at day E10.5 [32]. According to protein annotations, there are 3 P4HAs: P4HA1-3. However, P4HA3 does not appear to exhibit P4H activity [33]. Furthermore, our proteome investigation (Fig 1A) reports that P4HA3 is absent or to the least, incomparably poorly expressed, compared to P4HA1&2, in the present model, although at the mRNA it is found at considerable levels [27]. Altogether, the P4H activity should rely uniquely on P4HA1&2, which are expressed at comparable levels (Fig 1A).

**Fig. 1.**
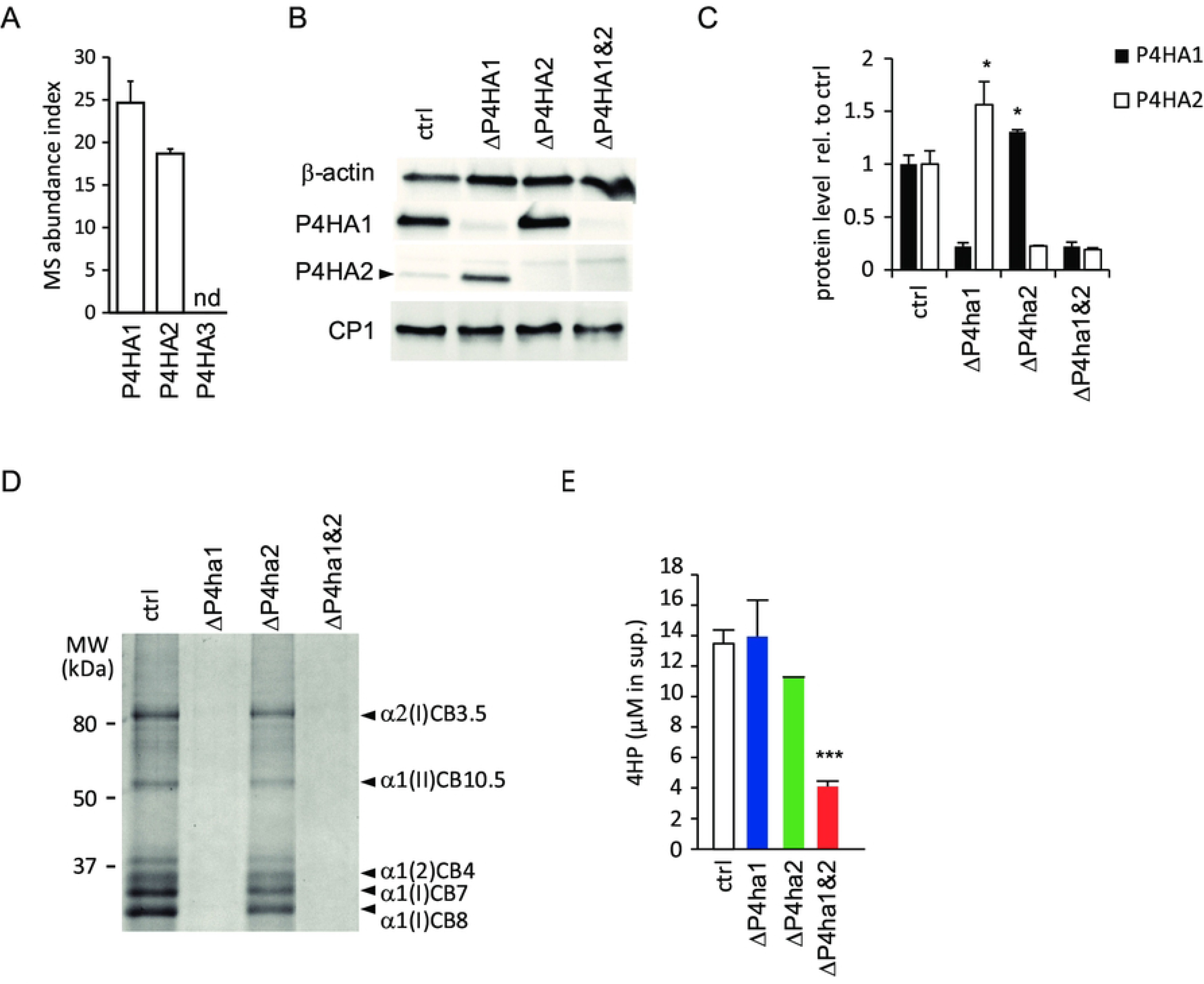
P4HA1 and P4HA2 exert quantitatively comparable but functionally distinct prolyl-4-hydroxylation in ATDC5 cells. (A) Mass spectrometry-based comparison of the expression levels of P4HA1, P4HA2 and P4HA3 in ATDC5 cells, based on the number of detected peptides in secretomes. nd: not detected. (B) Western blotting analysis of cellular fractions upon targeting of *P4ha1* and/or *P4ha2* genes using adenoviral transduction of CRISPR/Cas9 constructs; CP1: C-propeptide of type I collagen α1 chain. (C) Remnant P4HA1 and/or P4HA2 protein levels in experimental groups, measured by label-free relative quantitative mass spectrometry. (D) P4HA1 but not P4HA2 ablation abrogates crosslinked collagen synthesis, as reflected by the electrophoretic profile of CNBr digests, analyzed by SDS-PAGE coupled to Coomassie blue staining. CNBr fragments were annotated based on gel band tryptic fingerprinting by mass spectrometry. (E) 4-hydroxyproline levels in secretomes, as measured by UPLS-MS after total acid hydrolysis. Across the whole figure, histograms express the mean value + SD. * p-value < 0.05, *** p-value < 0.001, versus control.

Western blotting (Fig 1 B) confirms drastically decreased levels of P4HA1 and/or P4HA2 in samples infected with the corresponding viral constructs compared to controls. However, it needs to be clarified that, in the experimental setup used here, transformed cells are not subcloned, to avoid the degeneration of their collagen assembly ability. Therefore, experimental samples other than controls have a heterogeneous genotype. Indeed, according to our quantitative proteome analysis (Fig 1C), experimental groups in which *P4ha1* and/or *P4ha2* are targeted *(ΔP4ha1, ΔP4ha2* and *ΔP4ha1&2*) retain about 20 % of the normal enzyme levels. As the invalidation strategy relies on crispr/Cas9-mediated genomic deletions, it means that 80 % of the corresponding cell populations are knocked out for the genes of interest, while the remaining 20 % possess normal enzyme levels.

### P4HA1 is required for collagen assembly in ATDC5 cells

P4HA2 is dispensable for collagen assembly, as found earlier [31], but the lack of P4HA1 results in near extinction of the assembled collagen fraction (Fig 1D). By itself, this demonstrates distinct functions of P4HA1&2. As a technical consequence, collagen PTMs cannot be investigated from the assembled collagen fraction, as there is none in *ΔP4ha1* and *ΔP4ha1&2*. Therefore, in the follow-up of this study, PTMs are compared from the soluble secreted fraction, which contains similar amounts of procollagen species irrespective of the genotype as judged from type I procollagen C-propeptide secretion levels (Fig 1B).

When examined at the amino acid level, the content in 4HP of the secreted proteome obtained from *ΔP4ha1&2* falls down to about 25 % of controls (Fig. 1E). The remnant 4HP level in *ΔP4ha1&2* is attributable to cells that escaped the genome editing as explained above. In contrast, *ΔP4ha1* or *ΔP4ha2* cells exhibit a 4-hydroxyproline (4HP) level similar to controls; therefore, the two enzymes have similar, complementary contributions to the overall P4H activity, and the dependence of collagen assembly over P4HA1 (Fig 1D) does not rely on quantitative, but rather, qualitative differences in the substrates of the two enzymes.

### The type I collagen map of P4HA1&2 substrate residues

To better understand the substrate specificity of P4HA1&2, we investigated the dependence of each P4H site within type I collagen α chains, on P4HA1, P4HA2 or both of them, still from the same samples. To achieve this, we examined by high resolution mass spectrometry all identified post-translationally modified versions of each tryptic peptide obtained from both COL1A1 and COL1A2 polypeptides and compared their respective levels through a label-free relative quantitation approach, highlighting the positions on α chain precursors of prolyl residues that are substrates of either enzyme. The conclusions are summarized in Fig 2, covering the vast majority of the 212 potential P4H sites (defined by XPG tripeptides) and are explained in more details in the next paragraphs.

**Fig. 2.**
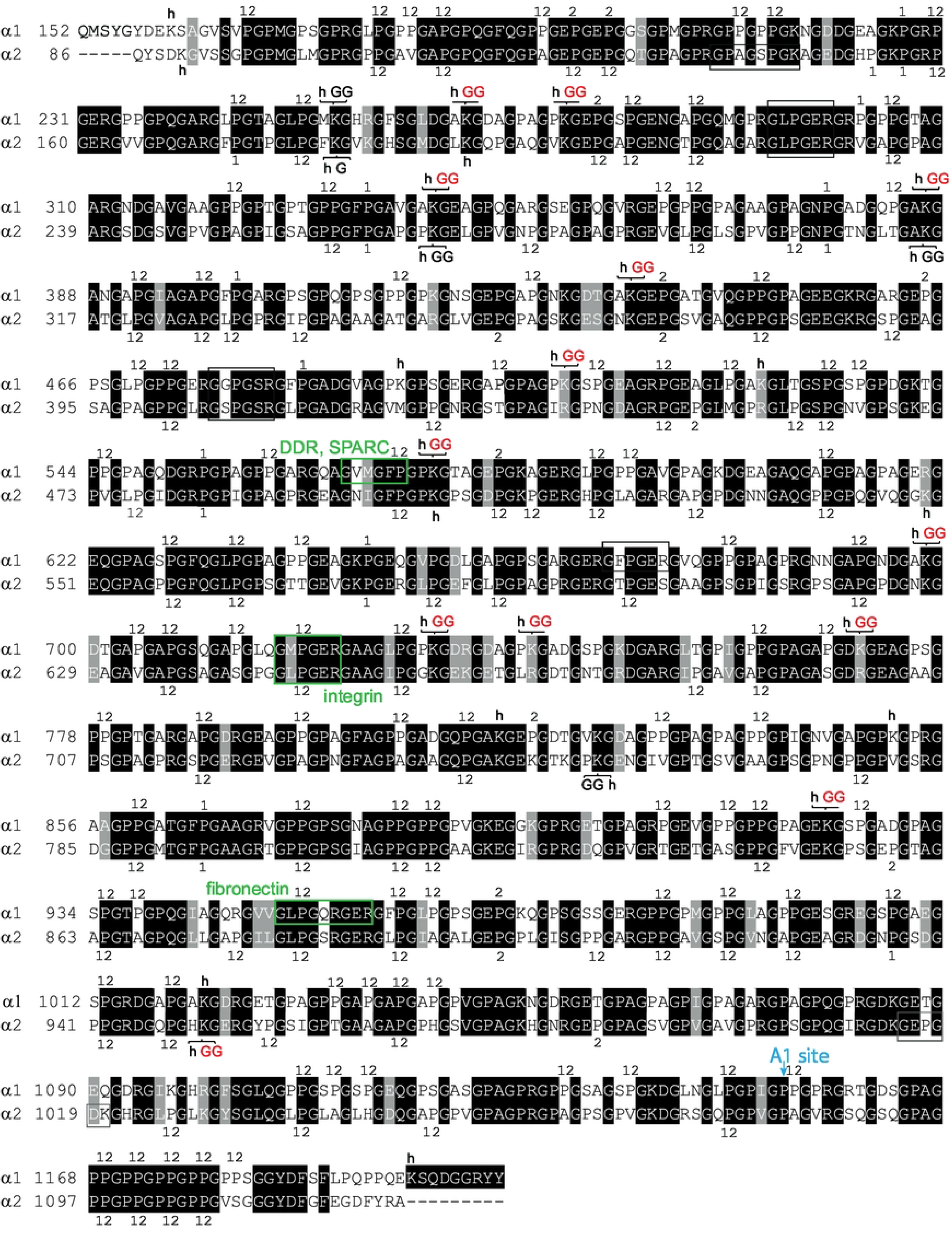
Mapping of the identified type I collagen post-translational modifications and their dependency on P4HAs. Residues are numbered according to their position in the precursors of α1 and α2 chains and annotated according to results of tryptic peptide characterization and relative quantification across the experimental groups by high resolution mass spectrometry. Strictly conserved residues are boxed in black. Functionally similar residues are boxed in grey. Regions boxed in light grey are not covered in the present analysis, most likely due to lack of chromatographic retention. Regions boxed in green were previously reported to bind functional partners according to annotations (reviewed in [34]). Residue annotations above and below the peptide sequences are as follows: 1, target of P4HA1; 2, target of P4HA2. Lysyl residues found modified are annotated with h, indicating optional (but major) hydroxylation and GG indicating optional glucosyl-galactosylation, in red when increased at least in *ΔP4ha1&2*. Of note, among the several GGs indicated in black, several could not be quantified due to interferences between glycosylation and tryptic cleavage at the same residue.

### P4HA1 and P4HA2 share most potential P4H sites as efficient substrates but exhibit individual substrate specificities at particular procollagen residue positions

A total of 160 sites are used as comparable substrates by both P4HA1&2, consistent with each enzyme being sufficient to achieve normal overall hydroxylation rates of procollagen (Fig 1E). Nevertheless, the two enzymes also exhibit respective substrate specificity, with 18 sites being targets of P4HA1 but not P4HA2 and *vice-versa* for 14 sites (Fig 2), as detailed below.

### Glutamic acid at the N-proximal position largely determines the substrate specificity of P4HA2

The invalidation of *P4ha2* was reported to cause decreased hydroxylation at most EPG tripeptides [27, 28]. The current data go along this conclusion (Fig 2), clarifying that 14 out of 20 characterized EPG sites are significantly better substrates of P4HA2 than of P4HA1, while 4 others are equivalent substrates of both and 2 are not found hydroxylated.

### P4HA1-dependent substrates are more heterogeneously defined by N-proximal residues.

In total, 18 residue positions are preferential substrates of P4HA1. Therefore, the numbers of specific substrates for each enzyme are in the same range and represent less than 10 % of the total potential 4HP sites. This is in agreement with the overall 4HP levels remaining similar to controls when either enzyme is missing (Fig 1E). The P4HA1-specific substrate sites have much more heterogenous N-proximal residues (summarized in Table 1) than those of P4HA2. In addition, each class of XPG tripeptide sequence (defined by the nature of the X residue) including P4HA1-specific substrate positions also includes positions accepted by both enzymes.

**Table 1.** N-proximal residue-dependence of P4HA1-specific substrates. The table summarizes the behavior of potential P4H sites found within tripeptide sequences FPG, KPG, NPG, RPG and HPG in terms of isoenzyme preference. P4H sites are identified here by their residue position within the corresponding α chain precursor polypeptide sequence. Of note, one FPG site (α1 671) is not characterized in the current data.

| Tripeptide sequence | P4HA1-specific | shared P4HA1/P4HA2 |
| --- | --- | --- |
| FPG | α1 335, 401, 485, 866<br>α2 174, 264, 795 | α1 572, 962<br>α2 501 |
| KPG | α1 227, 647<br>α2 156, 576 | α2 513 |
| NPG | α1 377<br>α2 306, 936 | α2 279 |
| RPG | α1 302, 554<br>α2 483 | α1 230, 518, 908<br>α2 447 |
| HPG | α2 153 | α2 519 |

### Collagen tripeptide analysis confirms the heterogeneity of P4HA1-dependent substrates in terms of N-proximal residues

To more directly question the influence of N-proximal residues over the substrate specificity of P4HAs, we employed a complementary strategy: The same samples were digested, this time with a recombinant collagenase [25], into GXY tripeptides and the hydroxylation rate of several GXP tripeptide sequences was then investigated by UPLC-MS (Fig 3). As opposed to the previous proteomics strategy relying on tryptic peptides, this approach provides a global, averaged overview of the hydroxylation rate of XPG sequences, independent on their position on the procollagen polypeptide or even on their α chain of origin. For instance, the 14 EPG sites found on both α1 and α2 chains of type I collagen, as well as those originating from other collagen types, depending on their relative levels, are in this analytical system considered as a whole from which the proportion that is prolyl-4-hydroxylated can be measured. It goes similarly, though with variable numbers of occurrences, for other XPG tripeptides, one limit being that the sensitivity of the method depends on the number of occurrences of a given tripeptide.

**Fig. 3.**
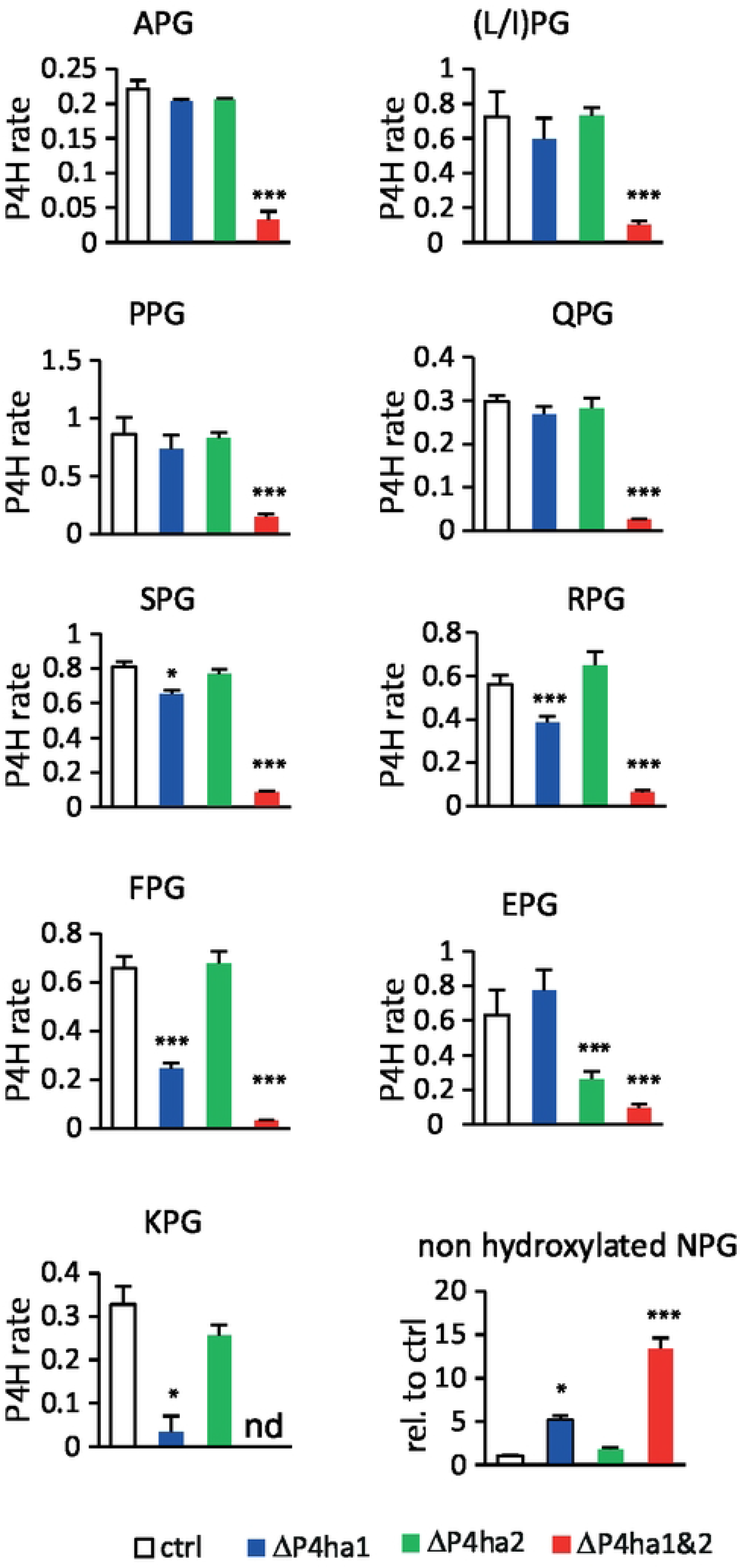
ATDC5 cell-secreted collagen tripeptide prolyl-4-hydroxylation analyzed by UPLC-MS. (A) Prolyl-4-hydroxylation rate for a subset of tripeptides of interest. (B) Relative levels of the non-hydroxylated state for another subset of tripeptides. Through the whole figure, histograms show the mean value, + SD. *: t-test p-value < 0.05. *** : t-test p-value < 0.001.

A first lesson from this investigation arises from the hydroxylation rate of individual XPGs in controls: while PPGs, L/IPGs (IPG and LPG are poorly distinguishable in the instrumental setup used here), SPGs, RPGs FPGs and EPGs reach hydroxylation rates from 60 to 80 %, APGs, KPGs and QPGs are in the 20-30% range, reporting a lower efficiency of P4HAs, collectively, towards these than the formers. At all investigated XPGs, *ΔP4ha1&2* exhibit radically dropped down hydroxylation levels compared to controls (Fig 3). However, APG, L/IPG, PPG, QPG and SPG tripeptides retain hydroxylation rates similar to controls in either *ΔP4ha1* or *ΔP4ha2*, (Fig 3), consistent with the sufficiency of either enzyme to achieve normal hydroxylation of these tripeptides. Hydroxylation of FPG and KPG, on one side, and EPG on the other side, drop down in *ΔP4ha1* and *ΔP4ha2*, respectively (Fig 3A), consistent with earlier reports [35] and the current findings at the tryptic peptide level (Fig 2). RPGs are significantly under-hydroxylated in *ΔP4ha1*, yet their relative levels remain above 2/3 of the controls, indicating a still fair ability of P4HA2 to hydroxylate these substrates. This is in agreement with our conclusions at individual sites as described in the previous paragraph.

For technical reasons, we were unable to measure hydroxylation levels of NPG tripeptides. Therefore, we estimated their hydroxylation based on the remnant levels of non-hydroxylated NPGs (Fig 3). These raise about 15-fold in *ΔP4ha1&2,* suggesting that the hydroxylation rate in controls must be in the highest range. They readily raise about 5-fold in *ΔP4ha1,* coherent with the notion that most NPGs are P4HA1-specific substrates (Fig 2).

To summarize, tripeptide analysis confirms at a global level the preferential hydroxylation of FPG, KPG, RPG and NPG tripeptides by P4HA1.

### P4HA activity is not required for prolyl-3-hydroxylation at the A1 site

P3H is believed to happen only at prolines standing on the X residue of GPP tripeptides where the proline in position Y is preliminarily 4-hydroxylated [17, 22]. Therefore, it should be dependent on P4HA activity. Our quantifications of total 3HP levels, measured from total hydrolysates (Fig 4A), confirm that *ΔP4ha1&2* exhibit reduced, but far from abrogated, 3HP levels. Among 3HP sites, the so-called A1 site is the only one being described as uniformly hydroxylated in animals, whatever the considered tissue [19, 21]. The tryptic peptide containing the A1 site also contains 2 potential 4HP sites, which we find are targets of both P4HA1&2 (Fig 2). However, in *ΔP4ha1&2*, this peptide retains a global hydroxylation rate (accounting for both 3HP and 4HP) slightly above 1 (Fig 4B), indicating that one residue remains hydroxylated independent on both P4HA1&2. Indeed, the main chemical state under which it is found (still in *ΔP4ha1&2*), carries a single hydroxylation, standing on the A1 site according to observed fragments (Fig 4C). Altogether, the current data indicate a more complex relationship between 4HP and 3HP than believed so far: although, in general, P3H is favored by P4HA activity (Fig 4A), at least at the A1 site, P3H is independent on P4H, occurring either at the neighboring residue or even anywhere else on the triple helix. We conclude that the mechanistic relationship between P4H and P3H is not as strict as previously thought, at least at particular sites.

**Fig. 4.**
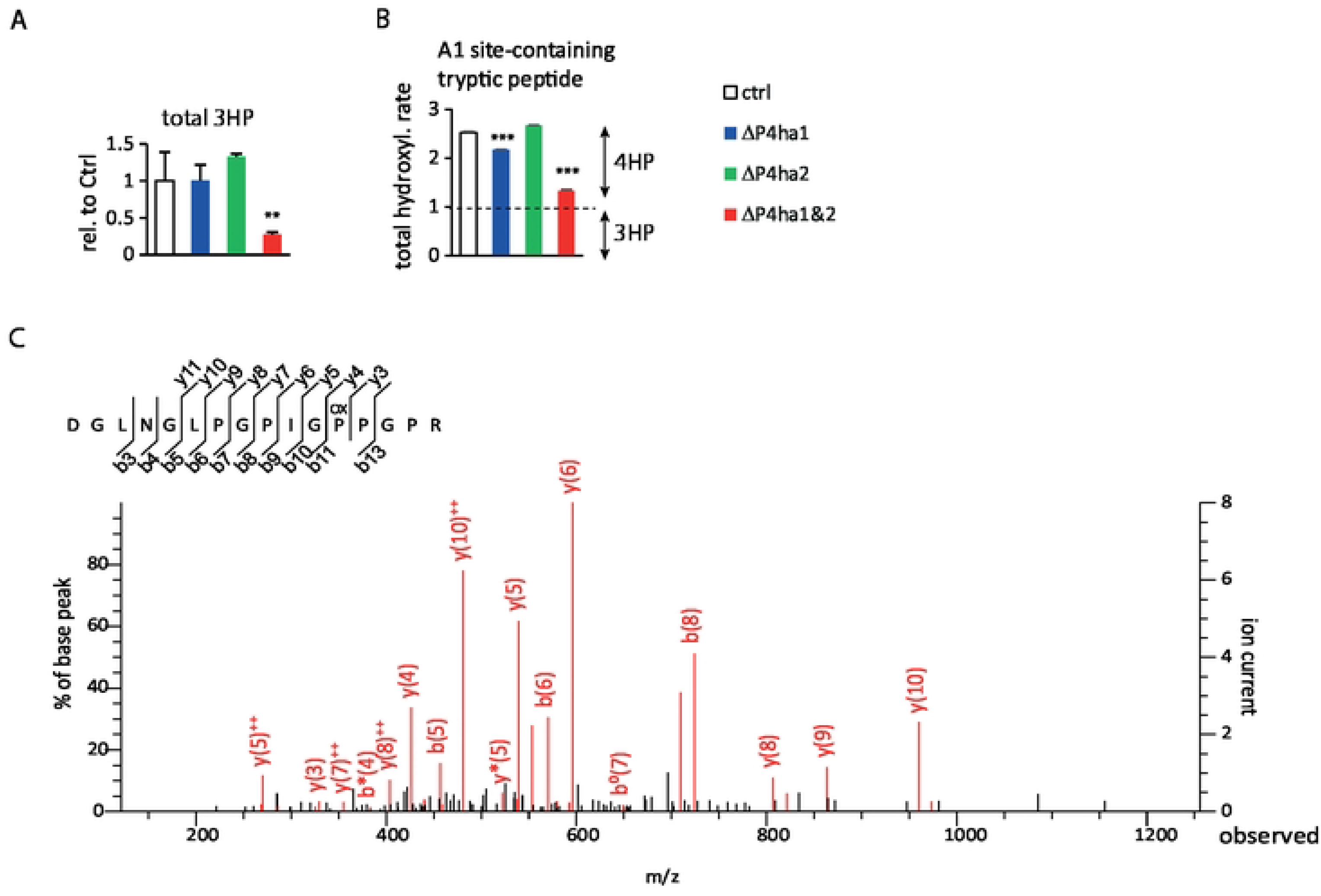
P4HA activity favors prolyl-3-hydroxylation overall but not at the A1 site, in ATDC5 cells. (A) Relative levels of 3-hydroxyproline in secretomes of ATDC5 cells as measured by UPLC-MS of total acid hydrolysates. (B) Total hydroxylation rate of the tryptic peptide containing the A1 site, as measured by LC-MS. (C) Fragmentation analysis of the tryptic peptide containing the A1 site, under its chemical state containing a single hydroxylation, as identified from the *ΔP4ha1&2* experimental group, showing that the target residue is the A1 site. Through the whole figure, histograms show the mean value, + SD. **: t-test p-value < 0.01. ***: t-test p-value < 0.001.

### P4HA activity restricts glucosyl-galactosylation of helical lysyl residues

In addition to prolyl residues, lysyl residues also undergo critical modifications, starting with hydroxylation on carbon 5, which can be extended with galactosylation and further glucosylation. These contribute to regulating the occurrence, valency and structure of collagen crosslinks [12–14] and possibly other aspects of collagen biochemical properties. In the present model, lysyl hydroxylation is evidenced at most potential sites within type I alpha chains (summarized in Fig 2), including the 2 N-telopeptide crosslinking sites, α1 K160 and α2 K90 and the unique C-telopeptide lysine, α1 K1197, as well as multiple helical sites. Remarkably, at all evidenced sites, occupancy by hydroxyl groups ranges from 50 to nearly 100 % in controls and remains stable in *ΔP4ha1&2*, indicating that lysl hydroxylation is independent on P4H. This is further supported by the similar hydroxylysine levels found in all groups after total alkaline hydrolysis (not shown). Galactosylation without further glucosylation is minimal (at best 5 %, in most cases undetectable, not shown). Two sites, α1 K254 (the N-proximal helical crosslinking site) and α2 K270, are mostly glucosyl-galactosylated in controls and their states are unaffected in *ΔP4ha1&2* (Fig 5A). A third site, α2 K753, is glucosyl-galactosylated at a rate around 1/3 independent on genotypes (Fig 5A). 13 sites exhibit increased glucosyl-galactosylation in *ΔP4ha1&2* to variable degrees. Glucosyl-galactosylation of α1 K275, α1 K437 and α1 K575 drastically increases in *ΔP4ha1&2* (Fig 5B), while at 10 other sites, glucosyl-galactosylation is increased in both *ΔP4ha1&2* and *ΔP4ha1* (Fig 5C). At 3 sites only, α1 K266, α1 K341 and α1 K770, *ΔP4ha2* is sufficient to induce, though more moderately, glucosyl-galactosylation (Fig 5C). The remaining sites exhibit low glycosylation in all genotypes and/or their glycosylation rates could not be calculated due to interferences of glycosylation at a given lysyl residue with subsequent tryptic cleavage at the same site (not shown). Coherent with the above, at the amino acid level after alkaline hydrolysis, glucosyl-galactosyl-hydroxylysine levels increase in *ΔP4ha1*, and more potently in *ΔP4ha1&2* (Fig 5D). Multiple lysyl hydroxylation/glycosylation sites, including some of these, were described *in vivo* previously [3, 6, 36, 37], suggesting their biological significance.

**Fig. 5.**
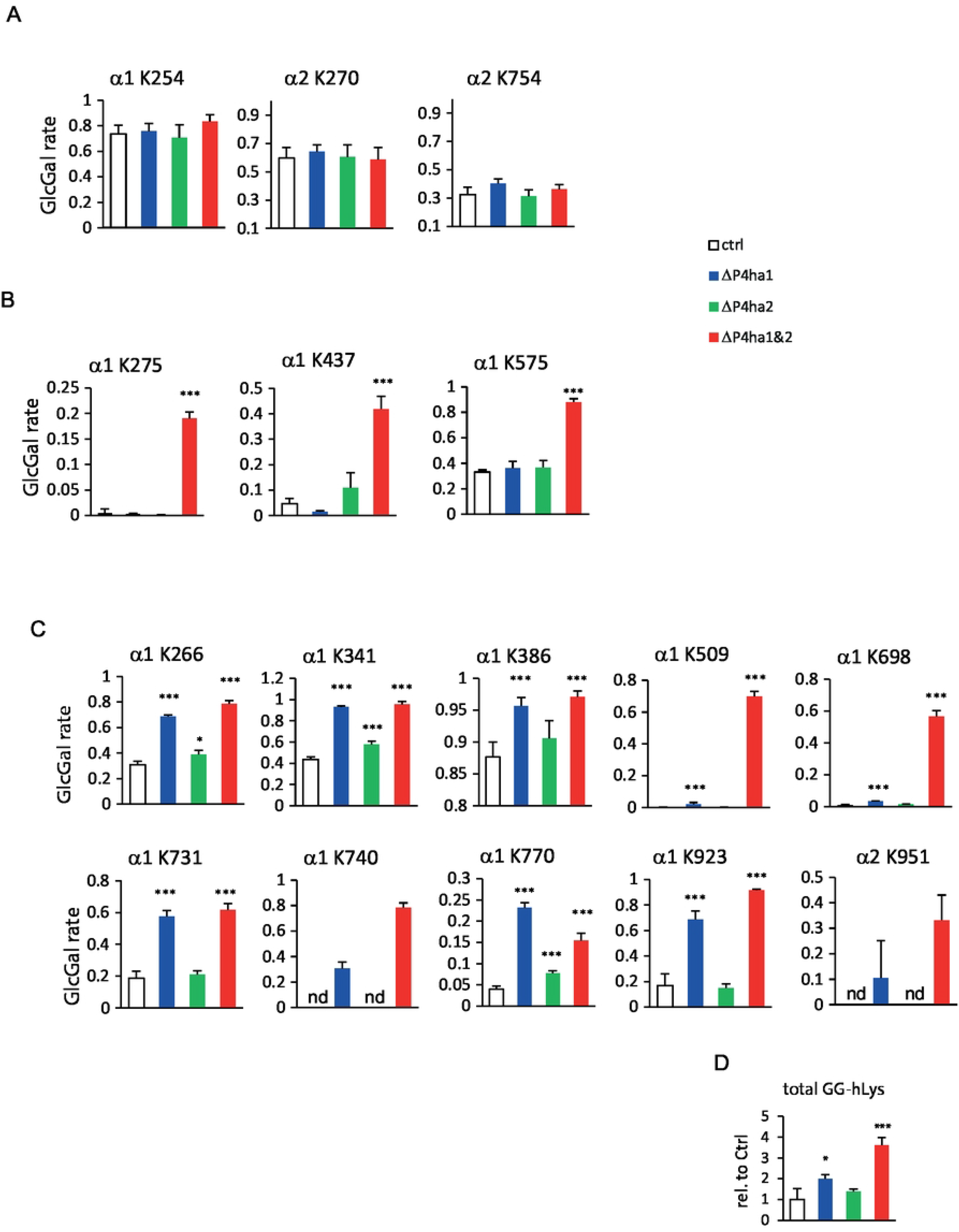
P4HA activity inhibits glycosylation at most potential sites of ATDC5 cell-secreted type I collagen. (A) Lack of regulation of glucosyl-galactosylation at 3 lysyl residue positions. (B) Glucosyl-galactosylation at 3 sites is induced only when both P4HA1 and P4HA2 are missing. (C) At 10 sites, the lack of P4HA1, and in 3 cases, the lack of P4HA2, is sufficient to induce glucosyl-galactosylation. (D) Relative levels of glucosyl-galatosyl-hydroxylysine in secretomes, as measured by UPLC-MS after total alkaline hydrolysis. Through the whole figure, histograms show the mean value, + SD. *: t-test p-value < 0.05. *** : t-test p-value < 0.001. nd: not detected. The numbering of residues refers to their position on α chain precursors.

Altogether, P4HAs have minimal influence over lysyl hydroxylation, but their lack results in induced glucosyl-galactosylation, with variable degrees depending on considered sites. P4HA1 is a stronger contributor to this regulation than P4HA2.

### Similar functions of P4HAs in mouse skin primary fibroblasts

Considering the complexity of the procollagen processing machinery, the consequences of P4HA functions could depend on the cellular environment, and therefore, the above findings may be relevant only to ATDC5 cells, the biological representativity of which is questionable. Therefore, the same experiments were performed in mouse skin primary fibroblasts. Results are similar in principles (Fig 6). P4HA1-2 are well detected in control cells but not P4HA3 (Fig 6A) and adenoviral constructs allow reduction of P4HA1 and/or P4HA2 in the same relative range (Fig 6B). Invalidating *P4ha1* drastically reduces mature collagen build-up (Fig 6C), as in ATDC5 cells. The levels of 4HP in secretomes are similar in controls, *ΔP4ha1* and *ΔP4ha2* but profoundly reduced in *ΔP4ha1&2* (Fig 6D). At the tryptic proteome level, the analytical depth is lower than in ATDC5 cells but interpretable data are in agreement with those reported in Fig 2 regarding the substrate specificity of P4HA1&2, although numerical values differ for many sites, along with basal hydroxylation levels in controls. At the tripeptide level also, conclusions are mostly similar, confirming EPG sites as P4HA2-specific and FPG, KPG and NPG sites, as P4HA1-specific (Fig. 6E-F), while other analyzable tripeptides are targets of both (not shown). Global 3HP levels (which are lower than in ATDC5 cells) are not significantly affected in *ΔP4ha1&2* (Fig 6G) and the peptide containing the A1 site retains a hydroxylation rate above 1 in *ΔP4ha1&2* (Fig 6H), suggesting that P3H is independent on P4H also in fibroblasts. Of note, however, in this model, MS/MS fragmentation did not allow ascertaining the residue position of remnant hydroxylation, due to lower signal. This model revealed itself less susceptible to lysyl glycosylation, as only 2 sites, α1 K575 and α1 K923 could be evidenced; yet, the two of them exhibit drastic increase in glucosyl-galactosylation, only in *ΔP4ha1&2* (Fig 6I).

**Fig. 6.**
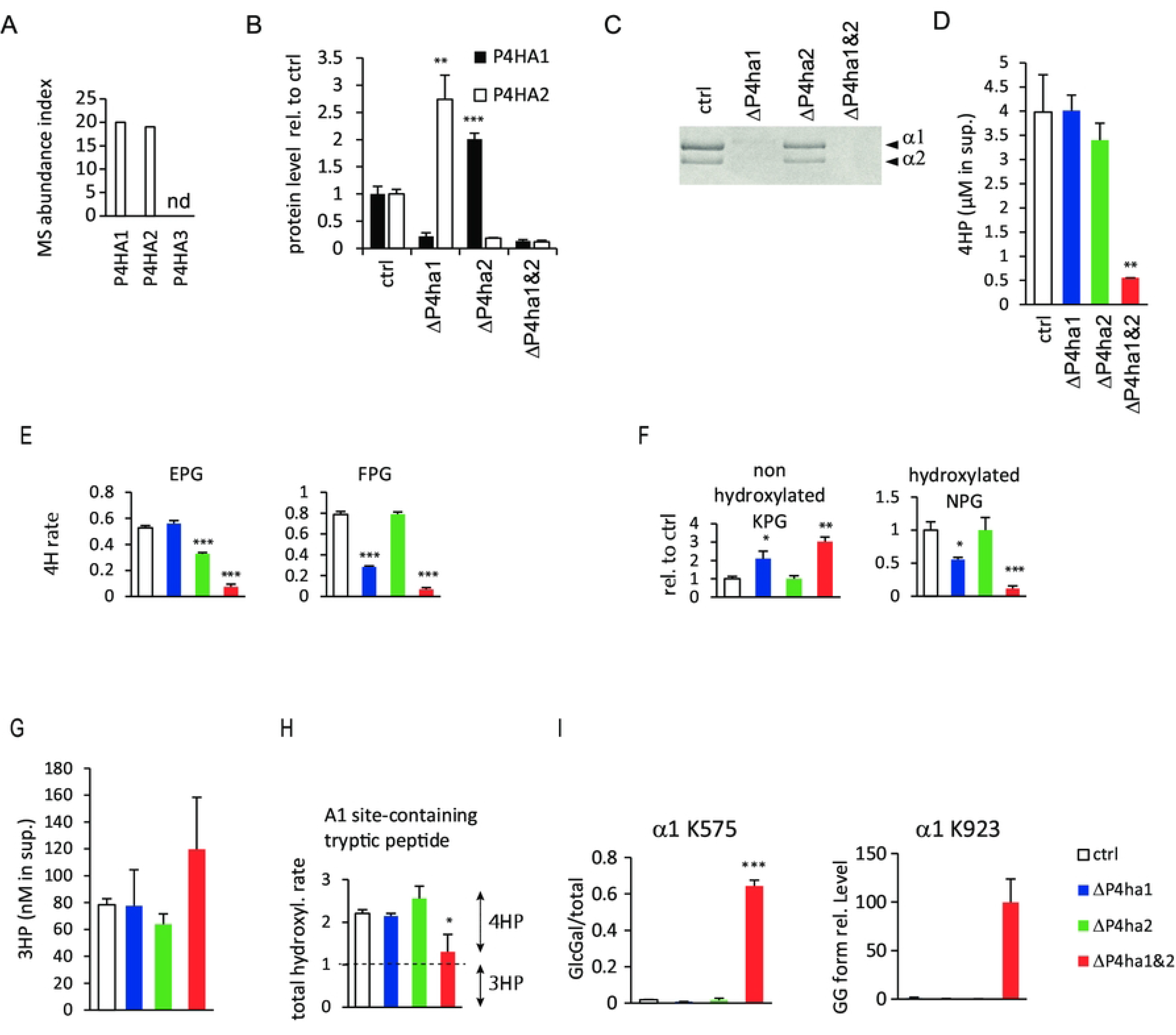
The consequences of P4HA1 and/or P4HA2 activity are conceptually similar in primary skin fibroblasts. (A) mass spectrometry-based comparison of the expression levels of P4HA1, P4HA2 and P4HA3 in fibroblasts, based on the number of detected peptides in secretomes. nd: not detected. (B) Remnant P4HA1 and/or P4HA2 protein levels in experimental groups, measured by label-free relative quantitative mass spectrometry. (C) P4HA1 but not P4HA2 ablation abrogates assembled collagen synthesis, as reflected by the electrophoretic profile of pepsin digests, analyzed by SDS-PAGE coupled to Coomassie blue staining. (D) 4-hydroxyproline levels in secretomes, as measured by UPLS-MS after total acid hydrolysis. (E) prolyl-4-hydroxylation rate for a subset of tripeptide exhibiting highest substrate specificity for either P4HA1 or P4HA2. (F) relative levels of either the non-hydroxylated state or the hydroxylated state for another subset of tripeptides also suggesting some degree of specificity towards P4HA1. (G) Relative levels of 3-hydroxyproline in secretomes of fibroblasts as measured by UPLC-MS of total acid hydrolysates. (H) total hydroxylation rate of the tryptic peptide containing the A1 site, as measured by LC-MS. (I) glucosyl-galactosylation rates at the 2 sites found modulated in fibroblasts. Across the whole figure, histograms express the mean value + SD. *: t-test p-value < 0.05, **:t-test p-value < 0.01, *** : t-test p-value < 0.001, versus control. The numbering of residues refers to their position on α chain precursors.

## DISCUSSION

### P4HA1 and P4HA2 share most potential P4H sites as substrates and each retains a limited set of enzyme-specific substrates

Collagen maturation involves incomparably high rates of PTMs that are crucial and/or determine details of its multistep assembly into polymeric, macromolecular networks. It is now increasingly clear that the PTMs are finely regulated in cells to control the biomechanical properties of the resulting collagen network. Until recently, P4HAs were not considered to contribute to this process, because 4HP is primarily required for triple-helix assembly, a prerequisite to collagen assembly. Recently, we and others started characterizing the substrate specificity of P4HA1 and P4HA2 *in cellulo* [27, 28] in models genetically invalidated for either enzyme-encoding gene [27, 28]. Here, for the first time, we characterize both enzymes in two independent models, using combinations of the two gene invalidations, including the double deficient one. It allows for identifying not only the collagen maturation sites that are significantly affected by the lack of either enzyme but also the ones at which there is partial compensation of one by the other. Our tryptic peptide analysis focuses on type I collagen because in the models used here, it is very predominant over other types and its PTMs were better studied in the past. Yet, the general conclusions drawn here for type I also seem to apply to types II, III and V in both models (not shown but available from the raw data), coherent with other recent conclusions [28]. Of note, our tripeptide analysis as well as the amino acid analysis reflect the average behavior of all collagen types, weighted by their relative abundancies. Either P4HA1 or P4HA2 alone is capable of achieving normal (or undetectably different from normal in the present experimental setup) overall 4HP level (Fig 1E) and the two share most potential P4H sites as substrates (Fig 2&3A). All residue positions found dependent on P4HA2 are part of EPG tripeptides (Fig 2) and this is confirmed at an overall level (Fig 3A&6E). Others found DPG tripeptides to also depend on P4HA2 [28], which would make sense in terms of functional residue classification, but in our hands, the single DPG tripeptide (α2 510) found in type I collagen α chains is hydroxylated by both enzymes, in agreement with our earlier finding [27] using a different mass spectrometry technology. Of note, we previously reported that EPG tripeptides, but also the single DPG tripeptide, were better hydroxylated in tendon than in bone and skin across distant species [27], suggesting functional consequences of the P4HA2-dependent hydroxylation, which will deserve further characterization.

Others previously [28] reported significant under-hydroxylation of collagen by P4HA1-deficient cells at positions following F, H, K, N, Q, R, S and with lower significance next to I, A and L residues. In our hands, P4HA1-specific sites are next to F, H, K, R and N (Fig 2, 3, 6E-F) but none of the others. In both models used here, the two enzymes are expressed at comparable levels (Fig 1A&6A) and the lack of either enzyme results in increased expression of the other (Fig 1B-C&6B), facilitating functional complementation. Also, here, both enzymes are characterized in the same models, allowing direct comparison of their abilities at hydroxylating a given position. At many positions, this demonstrates that the lack of both enzymes has a much stronger effect than the sole lack of P4HA1, leading us to define stringent criteria for considering one enzyme as specific (see the methods section). According to those criteria, the influence of N-proximal residues is insufficient to fully define the substrate specificity of P4HA1 as there also are FPG, HPG, KPG, RPG and NPG sites that are target of both enzymes or of none of them (Fig 2 & Table 1). The mechanisms dictating the specificity of P4HA1 remain to better characterize.

### Although quantitatively moderate, the respective substrate specificities of P4HA1 and P4HA2 have decisive functional consequences

The current study compares for the first time the influence of both enzymes on the macromolecular collagen assembly. Although each enzyme is capable on its own to achieve 4HP levels comparable to controls, P4HA1, but not P4HA2, is strictly required for collagen assembly (Fig 1D&6C), demonstrating the qualitative, in addition to quantitative, importance of P4H in collagen maturation. This contradicts previous findings, as P4HA1 was found required for type IV, but not for type I collagen assembly in early mouse embryogenesis and in cultured embryonic fibroblasts [32]. The discrepancy may be related to the distinct methods used in the previous and current studies to characterize collagen assembly. Still, the current finding reflects a fundamental difference between the 2 enzymes. Also, we trust that our findings at a molecular level reflect a dramatic loss of ECM stability in cultures lacking P4HA1, because the corresponding cell layers are incomparably more fragile (not shown), already at the beginning of the fractionation procedure. Of note, 4HP is required not only for triple helix stabilization but also for multiple collagen domains (reported in Fig. 2) to bind functional partners (reviewed in [34]); according to the present findings, all of these are targets of both enzymes and therefore their functionality should be retained when P4HA1 only is missing. Further studies will be required to clarify the requisite function of P4HA1 in collagen assembly as it is defined here.

### P4H is not systematically required for prolyl-3-hydroxylation to occur

Along with other recent work [30], our findings revisit previous consensual concepts in the mechanisms of collagen P3H. Although at the global level we confirm that P3H is favored by P4H in ATDC5 cells, we find that P3H at the A1 site (Fig 4A-B) is not. A previous work suggested that *P4ha1*(-/+), *P4ha2*(-/-) mouse skin exhibited increased P3H at the A1 site, compared to wild type littermates [29]. However, P3H levels of individual residue positions are difficult to quantify precisely due to the multiple PTM states under which the corresponding peptide sequence must have been found in this context. In our view, this issue may have biased the conclusions reported in [29], especially considering that the observed fold changes, though significant statistically, remained in a very close range (95 versus 96 % occupancy). Indeed, the A1 site is largely hydroxylated *in vivo* in any context [19, 21] and therefore, is not to be regulated. It is the main substrate of the P3H1 complex [19], involving particular chaperones [38–40] which could allow P3H1, unlike other prolyl-3-hydroxylases, to act before P4H, to ensure full P3H at this site. At this point and considering our confirmation that overall, the P3H level drops down in *ΔP4ha1&2* in the ATDC5 model (Fig 4A) but not in fibroblasts (Fig 6G), we conclude that the dependency of P3H over preliminary P4H is not as strict as previously consensually accepted [17, 22] and that P3H1 in particular escapes it, at least at the A1 site, which is its major substrate. The other way around, recent findings indicate that the lack of P3H1 increases P4H, particularly at P4HA2-dependent sites [30], as well as lysyl modification at several of the sites reported here. Altogether, the functional relationship between P4H and P3H turns out more complex and intricate than previously thought.

### P4H inhibits lysyl glycosylation of procollagen

Lysyl hydroxylation is not found modulated by P4HAs as judged from tryptic peptide analysis (Fig 2) and from amino acid analysis after alkaline hydrolysis (not shown), but the further maturation of hydroxylysine by glucosyl-galactosylation, is fully inhibited by P4HAs at 13 of 16 potential sites in ATDC5 cells (Fig 5). Therefore, lysyl hydroxylation is independent on P4H while glucosyl-galactosylation must occur before massive P4H. Remarkably, in fibroblasts, the number of characterized potential sites drops down to 2 but glucosyl-galactosylation remains inhibited at both of them by P4HAs (Fig 6I). Confronting results obtained in both models suggests that each cell type has a glycosylation machinery able to act at intrinsically defined lysyl sites, while the rate at which it acts is modulated by P4H. In agreement, the collagen-specific chaperone Hsp47 inhibits P4HA activity and increases lysine modifications [5, 41, 42]. When P4HA1 alone is missing, the overall P4H level of secreted procollagen is normal (Fig 1E) and yet glucosyl-galactosylation increases overall (Fig 5D) and at particular sites (Fig 5C). The influence of P4HA2 is much weaker, as 3 sites only exhibit significantly increased glucosyl-galactosylation when it is missing (Fig 5C). Of note, 4 sites reported here were previously found significantly more glucosyl-galactosylated in *P4ha1*(-/+), *P4ha2*(-/-) mouse skin [29], confirming *in vivo* the pertinence of the herein more exhaustive findings. Altogether, the substrate specificity of P4HA1 and P4HA2, combined with their extrinsically controlled expression and activities, could not only establish tissue-specific P4H profiles but also orchestrate the finely-tuned P3H and lysyl hydroxylation/glucosyl-galactosylation patterns. Further studies will be required to better delineate the mechanisms involved and the downstream functional consequences.

## METHODS

### ATDC5 Cell culture and genome edition

The ATDC5 model was used strictly as described in [27], except that two rounds of adenoviral infections were performed (to create double mutants). In the first round, cells were infected at a MOI of 100 with either a single-guided adenoviral CRISPR/Cas9 construct targeting human *Hexose-6-Phosphate Dehydrogenase* (gRNA sequence: CACCTGCGAAGGTCGGCGTC, used here as negative control) or with one targeting *P4ha2* (gRNA sequence: AGATCAGCTGCCGACCCCGA). Cells were allowed to reach confluency, were passaged 1:6 into new 12-well plates and 48 hours later, infected again with either the construct targeting *H6pd* or a novel one targeting *P4ha1*, expressing two gRNAs (gRNA1 sequence: CCAGGATGAACTTAGGATTC; gRNA2 sequence: CCATGACATCATCTCTGACG). Adenoviral constructs were provided by Vectorbuilder. Cells were allowed to reach confluency, the medium being changed every day, before insulin stimulation. 3 independent infections were used per experimental group as specified. Differentiation was stimulated for 16 days in all experiments and the medium was supplemented with 50 µM ascorbate, the medium being serum-deprived over the last 2 days. These experiments were repeated twice independently.

### Mouse primary skin fibroblast extraction and cell culture

Mouse skin fibroblasts were isolated from 27 to 37-day-old wild type male CD57BL6/J mice sacrificed at our local facility in the context of other studies, as described in [43]. At passage 2, cells were cultured in DMEM/F12 medium supplemented with 20 % FCS. For adenoviral infections, cells were plated at 15,000 cells/cm² in 12-well culture plates and grown as monolayer to 80 % confluence, then, they were infected with the same constructs as for ATDC5 cells, but all infections were performed in a single round, each virus at a MOI of 50. Three independent infections were used per experimental group. Cells were grown to confluency and collagen production was stimulated for 6 days with 500 µM ascorbate and 50 µM proline, the medium being serum-deprived over the last 2 days. These experiments were repeated twice independently.

### Protein fractionation from ATDC5 cells, SDS-PAGE, western blotting analysis, Coomassie staining and sample preparation for LC-MS

These aspects were managed strictly as described in details in [27, 31].

### Protein fractionation and sample preparation from primary fibroblasts

Secretomes were collected and prepared strictly as for ATDC5 cells. From this point all steps were performed at 4°C. Cell layers were collected in 1 ml PBS containing 1 % Triton X-100, vortexed and incubated on wheel for one hour and centrifuged at 15 000 g for 5 min. The soluble fraction containing most cell components was collected and the pellet was rinsed once again in the same buffer and 3 times in water. Further extraction was performed in 1 ml acetic acid 0.5 M for 3 days. This fraction contained minimal collagen. Further pepsin extraction was performed in the same buffer containing 10 µg pepsin for 3 days, after what the pellet was negligible. Pepsin extracts were precipitated with DOC/TCA, resuspended in Laemmli Buffer and equal fractions were loaded onto SDS-PAGE prior to Coomassie staining. Sample preparation for LC-MS was strictly as described for ATDC5 cells.

### LC-MS analysis

Samples were resuspended in 100 µl 2 % ACN, 0.1% formic acid (FA), clarified upon centrifugation and 75 µl were transferred to sampler tubes. Analysis was performed on a timsTOF Pro 2 mass spectrometer coupled with a nanoElute 2 liquid chromatography system (Bruker Daltonik GmbH, Germany). 5 µl aliquots were injected onto a desalting column (C18, 5 mm x 0.3 mm, 5 µm particle size, Thermo) and fractioned on a 150 µm i.d. x 100 mm, 1.9 µm particle size, 150 A° pore size, Bruker column, using a two-column separation method using a 60 min, 2-30 % acetonitrile gradient at a flow rate of 0.5 μL/min. The column temperature was set to 50 °C. Peptides were introduced into the mass spectrometer via a CaptiveSpray nanoelectrospray ion source (Bruker Daltonik GmbH) through a 20 µm emitter at an electrospray voltage of 1.6 kV, an ion source temperature of 180 °C, and a dry gas of 3 L/min. Data collection was carried out using DDA-PASEF as follows: A total of 10 PASEF ramps targeted precursors with a charge between 2 and 5 and a cycle time of 1.17 s. Each PASEF ramp was set to a ramp time of 100 ms. Ion mobility and mass were calibrated for the mobility range 0.6–1.60 V·s cm^−2^ (1/K0) using Agilent Tune mix as a reference. The collisional energy increased linearly from 20.0 eV at 0.60 V·s cm^−2^ (1/K0) to 59.0 eV at 1.60 V·s cm^−2^ (1/K0). Both MS and MS/MS spectra were recorded within a scan range of 100–1700 m/z. TIMS accumulation time was set to 100 ms, with a target precursor intensity of 20,000 arbitrary units (au) and a minimum threshold of 2500 au. Active exclusion was set to 0.4 min. Absolute intensity thresholds were set to 10 for mass spectra peak detection and 5000 for mobilogram peak detection.

### Peptide and PTM identification, including curation of residue localization of PTMs

Raw data (.d folders) were processed with Mascot Distiller bound to Mascot Server (both from Matrix Science Ltd) allowing in the setup up to 8 PTMs per peptide. Identification was managed by interrogation of the Swissprot mouse database, allowing 10 ppm deviation of MS data and 0.1 Da deviation of MS/MS data, with trypsin cut, one allowed miss-cleavage and considering proline hydroxylation, lysine hydroxylation, lysine galactosylation, and lysine glucosyl-galactosylation, together with methionine oxidation, with a threshold FDR at 1 %. All PTMs of interest were inspected manually based on the fragments automatically selected during Mascot interrogation, to clarify the residue positions of PTMs based on fragmentation, when possible. P3H and P4H were distinguished strictly based on their positioning at X versus Y residues of XYG tripeptides, respectively.

### Relative quantification of individual molecular states

The peptide tables (exported from Mascot distiller) were exported to an Excel file and processed using a macro to create one file per peptide sequence, combining all PTM and charge states as well as the corresponding quantitative data based on peak areas, calculating, based on this, the average hydroxylation, galactosylation and glucosyl-galacosylation rates, expressed as average + SD, together with t-test p-values for each experimental group versus controls and highlighting significant changes. MS/MS data corresponding to peptides exhibiting significant changes were manually inspected to clarify whether or not the fragments discriminated the positions of the corresponding PTMs.

### Total hydrolysis and UPLC MS amino acid analysis

400 µl aliquots of supernatants were transferred to Teflon-capped thermoresistant tubes, ethanol-precipitated, dried and equilibrated in a globe box kept under constant nitrogen flow until the ambient oxygen became undetectable. They were then resuspended in 100 µl HCl, 6 N, for proline analysis, or NaOH, 2 N, for lysine analysis, tightly closed, heated to 110 °C overnight, neutralized with either 6 N NaOH or 6 N HCl, and clarified by centrifugation. 10-50 µl (depending on the amino acid to investigate) aliquots were used for AccQ-Tag reaction [44] as recommended by the supplier (Waters). 2-20 µl (depending on samples and considered amino acids) samples were analyzed on a Acquity Premier UPLC system equipped with a Quaternary Solvent Manager, a Flow-Through Needle manager injector, a column manager oven and interfaced to a UV detector (for quality control only) and a single quadrupole detector (QDa). Chromatographic separation was performed on a CORTECS® C18 (1.6 µm, 2.1 × 50 mm) column with a CORTECS® C18 VanGuard™ pre-Column (1.6 µm, 2.1 × 15 mm) at 55 °C. The gradient elution at 0.5 ml/min flow rate was 1-25 % ACN in water with 0.1 % FA, over 5 min. The QDa mass detector was operated in the positive ion mode using selected ion recording (SIR) at 10 V cone voltage. The source temperature was set to 120 °C and the nitrogen flow was kept at 1200 L/h during the whole analytical procedure. Proline, hydroxyproline, lysine, hydroxylysine, galactosyl-hydroxylysine and glucosyl-galactosyl-hydroxylysine were monitored at m/z = 286.2, 302.2, 244.1, 252.1, 333.45 and 413.45, respectively. Standards consisted in mixtures of free amino acids including proline, 4-hydroxyproline, 3-hydroxyproline (the two have well separated retention times), lysine and hydroxylysine in the 10^-10^ to 2.10^-8^ mol range, similarly processed through total hydrolysis and AccQ-Tag reaction. Acquisition and data processing, including quantification based on calibration curves, were processed with Empower software (all from Waters).

### Collagen tripeptide analysis by UPLC

The method was derived from [25]. 1 ml clarified culture supernatant was DOC/TCA-precipitated, resuspended in 100 µl collagenase buffer (20 mM Hepes pH 7.5, 30 mM NaCl, 2 mM CaCl2), heated to 95°C for 5 min, quick-chilled, and digested with 200 ng Brightase C (reprocell) overnight at 37°C. 40 µl aliquots were used for AccQ-Tag reaction and UPLC measurement as described above. Calibrants consisted in synthetic peptides with the following sequences: GLPGPPGAPGPQGFQGPPGEPGEPGGSGPMGPRGPPGPP and GLOGPOGAOGPQGFQGPOGEOGEOGGSGPMGPRGPOGPO (where O stands for 4HP), and served to ascertain the retention times of the corresponding tripeptides. Hydroxylation rates were calculated based on the absolute peak areas for the hydroxylated and non-hydroxylated states of each tripeptide sequence. In cases where either one could not be measured, the relative intensity of the other was used as marker of hydroxylation.

### Thresholds and statistics

All experiments were performed twice independently (at different times), with parallel independent experimental triplicates. The numeric data reported here correspond to one experiment per model, but are in conceptual agreement with the other experiment. In proteome analysis, a given hydroxylation site was considered dependent on a given enzyme only if the P4H rate in the corresponding mutant was statistically different from the control and closer, in absolute values, to the level observed in the double mutant than in the controls. Values are expressed as mean + SD. The significance of group-to-group comparisons is estimated based on unbiased student t-tests.

## Data availability

The mass spectrometry proteomics data have been deposited to the ProteomeXchange Consortium via the PRIDE partner repository with the following dataset identifiers: for data obtained from ATDC5 cells, PXD075804, for primary fibroblasts, PXD083268.

## Declaration of AI use

During preparation of this manuscript, the authors used Claude from Anthropic, strictly to check the main document for consistency with the figures and to obtain suggestions for improving the clarity and phrasing of the text. All suggestions were reviewed and edited by the authors, who take full responsibility for the content of the publication.

## Data availability

The mass spectrometry proteomics data have been deposited to the ProteomeXchange Consortium via the PRIDE partner repository with the following dataset identifiers: for data obtained from ATDC5 cells, PXD075804, with reviewer password G90QtR6tMxKq and for primary fibroblasts, PXD083268, with reviewer password dUeu3wjO8ae6.

## ABBREVIATIONS

3HP: 3-hydroxyproline
4HP: 4-hydroxyproline
COL1A1: α1 chain of type I collagen
COL1A2: α2 chain of type I collagen
CNBr: cyanogen bromide
ΔP4ha1: P4ha1-deficient cells
ΔP4ha2: P4ha2-deficient cells
ΔP4ha1-ΔP4ha2: P4ha1-P4ha2-deficient cells
LH: Lysyl Hydroxylase
P3H: Prolyl-3-Hydroxylation
P4H: Prolyl-4-Hydroxylation
P4HA: Prolyl-4-Hydroxylase Alpha subunit
PTM: post-translational modification
EIC: Extracted Ion Chromatogram

